# Validation and multi-site deployment of a lyophilized qRT-PCR reagent for the molecular diagnosis of avian influenza and rabies in Sub-Saharan African regions

**DOI:** 10.1101/2025.01.16.633398

**Authors:** Petra Drzewnioková, Irene Brian, Marzia Mancin, Andrea Fortin, Morgane Gourlaouen, Angélique Angot, Mamadou Niang, Isaac Dah, Kouramoudou Berete, Adama Diakite, Fatou Tall Lo, Clement Meseko, Emilie Go-Maro, Valeria D’Amico, Viviana Valastro, Baba Soumare, Paola De Benedictis, Isabella Monne, Valentina Panzarin

## Abstract

Molecular methods are widely accepted as gold standard techniques for the laboratory diagnosis of most human and animal pathogens. However, most molecular protocols rely on reagents that need to be transported and stored at a freezing temperature, a requirement that might affect their reliability in areas where the cold chain cannot be guaranteed. Over the years, several lyophilized molecular products have been marketed to circumvent this issue. We therefore evaluated the feasibility of replacing liquid reagents with freeze-dried formulations for the molecular diagnosis of avian influenza (AIV) and rabies (RABV) viruses, two priority zoonotic pathogens widely spread in Sub-Saharan Africa. Among the available kits, we selected the Qscript lyo 1-step kit (Quantabio) due to its easy-to-use features, single-reaction format, and preliminary performance assessment. Through a more in-depth evaluation, we determined its analytical and diagnostic performance and formulation stability, and obtained results comparable to those of standard liquid master mixes. Notably, for the detection of divergent lyssaviruses, the lyophilized reagent’s sensitivity was affected by suboptimal complementarity between the oligonucleotides and the target sequences. Finally, a multi-site evaluation in four veterinary diagnostic laboratories located in Sub-Saharan Africa demonstrated the successful deployment of AIV and RABV assays utilizing freeze-dried reagents, which can interchangeably replace liquid master mixes. Altogether our results indicate that the Qscript lyo 1-step kit (Quantabio) represents a valid alternative to wet reagents for the molecular diagnosis of avian influenza and rabies, and has the potential for broader applications to other relevant infectious diseases upon proper validation.

**Author summary:** Molecular diagnostic protocols rely on reagents that need to be transported and stored at freezing temperatures. Meeting this requirement can be challenging in areas where the maintenance of the cold chain is not guaranteed, such as in sub-Saharan Africa. Our study aimed to assess the feasibility of using lyophilized reagents as a replacement for liquid reagents in the molecular diagnosis of two widespread zoonotic pathogens in Sub-Saharan Africa, namely avian influenza and rabies. To accomplish this, we selected a commercially available lyophilized reagent based on its format and performance characteristics. We conducted a laboratory validation to assess the use of the lyophilized reagent throughout the entire diagnostic process. We also conducted a reproducibility test involving African laboratories as potential end-users. Our findings confirm that the lyophilized reagent can replace traditional liquid reagents to diagnose rabies and avian influenza and suggest its possible use for a wider range of infectious diseases after undergoing appropriate validation.

## Introduction

The strong connections between the health of humans, animals and the ecosystems recognized by the One Health holistic approach has led to the establishment of the Global Plan of Action for One Health (GPA) shared within the quadripartite alliance [1,2] among the Food and Agriculture Organization of the United Nations (FAO), the UN Environment Programme (UNEP), the World Health Organization (WHO) and the World Organization for Animal Health (WOAH). The GPA identifies as top priorities not only emerging zoonotic diseases with epidemic and pandemic potential, but also neglected and endemic diseases of zoonotic and vector-borne origin [2]. Indeed, while the disruptive impact of epidemics and pandemics caused by the emergence of zoonotic pathogens appears obvious, the indirect consequences in terms of missed opportunities for the control, prevention and treatment of endemic and neglected diseases should not be overlooked [3–5]. In this view, zoonotic influenza (including avian influenza, AI) and rabies have been long recognized as potential showcases for the One Health inter-sectorial collaboration [6–8].

AI is a highly contagious disease caused by influenza A viruses (AIV; genus *Alphainfluenzavirus,* family *Orthomyxoviridae*), which can be classified based on their levels of pathogenicity to chickens or the composition of the haemagglutinin (HA) cleavage site, resulting in designations as either low pathogenicity avian influenza (LPAI) or high pathogenicity avian influenza (HPAI). So far, only viruses of the H5 and the H7 subtype have proved to be able to evolve into a HPAI form, displaying a heightened ability to cause severe disease and death in infected poultry. Among the multiple subtypes of AIV reported so far in domestic birds, LPAI H9N2 and HPAI H5N1 viruses are currently the most widespread in the poultry population [9]. Both subtypes can cause significant economic losses to the poultry industry, being responsible for a remarkable number of infections in mammals and for sporadic spillover events in humans [10,11]. These viral threats have a particularly devastating impact in developing countries, such as the Sub-Saharan African territories, where poultry farming is a highly growing sector counting on approximately 1.1 billion of poultries [12] and, at the same time, an important source of income for several traders and rural households [13]. Indeed, HPAI H5N1 viruses of the A/Goose/Guangdong/1/1996 (Gs/GD) lineage [14] and G1-like lineage H9N2 viruses are now entrenched across the poultry population of multiple African countries. Combined with poor surveillance, limited response capacity and deficient reporting, this complex epidemiological scenario creates the opportunity for virus evolution and paves the way for the emergence of strains with unexpected zoonotic potential [15,16].

Rabies, an ancient although preventable disease, still kills an estimate of 60,000 people every year [6] and mostly affects the poorest communities in rural Africa and Asia [17–19]. Rabies can also occasionally result on livestock’s death due to spillover events, determining a 6% income loss of the 8.6 billion USD economic losses per year [20] that is expected to impact on the fragile economies of endemic countries [17,21]. Despite rabies is potentially caused by all members belonging to *Lyssavirus* genus (family *Rhabdoviridae*), rabies virus (RABV) is the most widely spread and the main responsible for human deaths worldwide. Indeed, the remaining 17 non-RABV lyssavirus species recognized to date [22] display restricted geographical and ecological niches, mostly remaining confined to their natural hosts. In Africa, six non-RABV lyssaviruses have been detected so far, namely Duvenhage virus (DUVV) [23], Lagos Bat Virus (LBV) [24], Mokola virus (MOKV) [25], Shimoni bat virus (SHBV) [26], Ikoma virus (IKOV) [27] and the recently discovered Matlo Bat Virus (MTLBV) [28], with rare spillover infections identified in domestic animals and humans [29]. Considering that infections with divergent lyssaviruses are able to cause severe clinical signs and impact on public health, international organizations recommend the use of a broad range of laboratory methods to diagnose both RABV and non-RABV lyssaviruses from animal suspect cases [30,31]. Nevertheless, rabies is maintained by a vicious circle of neglect, in which poor surveillance and reporting lead to underestimating the burden of the disease and, consequently, to making insufficient efforts for its control and prevention [6,12,13,17,21]. Thus, improved rabies surveillance and diagnosis are key drivers for human and animal health and for measuring the impact of control programs, ultimately offering reliable data to decision-makers [17,32,33].

In the last decades we have witnessed the massive implementation of molecular techniques that have greatly contributed to the success of surveillance and control programs for infectious diseases [34]. Regulatory health authorities have long recognized the use of molecular tests for AIV detection and typing, and largely rely on the rapidity of such methods for the prompt confirmation and notification of outbreaks [35,36]. On the opposite, rabies molecular testing techniques have only recently been included in the WOAH and WHO diagnostic manuals [30,31], albeit their suitability has been widely demonstrated by direct comparison with gold standard methods [37,38]. So far, high-income countries have been the main beneficiaries of such fast and accurate molecular methods [17,21,39] as their settings can guarantee cold-chain maintenance for proper transportation and storage of reagents and samples [40]. Unfortunately, this basic requirement is a struggle in low- and middle-income countries of tropical and subtropical areas, whose operability is often prevented by inadequate investments and unfavorable climate conditions [41] despite the technical skills achieved by the laboratory personnel through consolidated cooperation programs [42,43]. Actually, we are aware of the difficulty in maintaining the recommended temperature for storing samples after delivery to diagnostic laboratories, and we well understand that such an inappropriate storage of samples and reagents at laboratory level is mostly due to the discontinuity of electricity supply. Altogether, these factors lead to the limited submission of samples for testing, which in turn reduces the surveillance capacity in the poorest countries [41]. In such contexts, the availability of diagnostic solutions that require only basic storage conditions would be highly beneficial. Great efforts have been made to develop and commercialize this kind of products for AIV and RABV detection, which are however mostly based on point-of-care (POC) antigenic tests whose sensitivity is lower than PCR-based methods [44–50]. An attractive alternative is represented by freeze-dried reagents for PCR-based applications, which promise to offer the sensitivity of molecular methods with less-strict requirements for reagents preservation [51]. While the application of these reagents is still unexplored for RABV detection, a number of independent studies have preliminary demonstrated the suitability of lyophilized formulations for the molecular diagnosis of avian influenza [44,52–56]. However, these studies neither assessed the performance of the lyophilized reagents in the actual conditions faced by low-income countries nor were they restricted to the evaluation of screening methods only, at the same time neglecting downstream subtyping assays. To explore more in depth the potentiality of freeze-dried amplification kits with basic storage requirements, we compared a commercial lyophilized reagent to wet master mixes employing real-time RT-PCR (qRT-PCR) and end-point RT-PCR methods for the detection and typing of avian influenza and rabies viruses. The suitability of the lyophilized reagent in replacing current wet master mixes was further assessed by determining its actual thermostability under stressful environmental conditions, as well as through an inter-laboratory reproducibility study that involved four Central Veterinary Laboratories (CVLs) located in Sub-Saharan Africa.

## Materials and Methods

### Selection of freeze-dried kits and molecular protocols

We performed an inventory of commercial freeze-dried master mixes meeting the following criteria: i) one-step RT-PCR/qRT-PCR for the detection of RNA targets; ii) suitability for TaqMan probes-based qRT-PCR assays; iii) adaptability to custom oligonucleotides. The performance of the selected product was extensively compared to standard liquid reagents employed in reference qRT-PCR and end-point RT-PCR methods for AIV and RABV detection and typing (Table 1) [30,31,36,57–65] the majority of which have been widely employed also in the CVLs taking part in this study. For analytical sensitivity, repeatability and thermostability tests, viral RNA was purified from viral isolates using the QIAamp Viral RNA kit (Qiagen). For diagnostic sensitivity and inter-laboratory reproducibility the NucleoSpin RNA kit (Macherey-Nagel) was employed. When performing the protocols described in [59,63], six microliters of the exogenous control intype IC-RNA (Indical) were added to the lysate during RNA isolation.

**Table 1.**
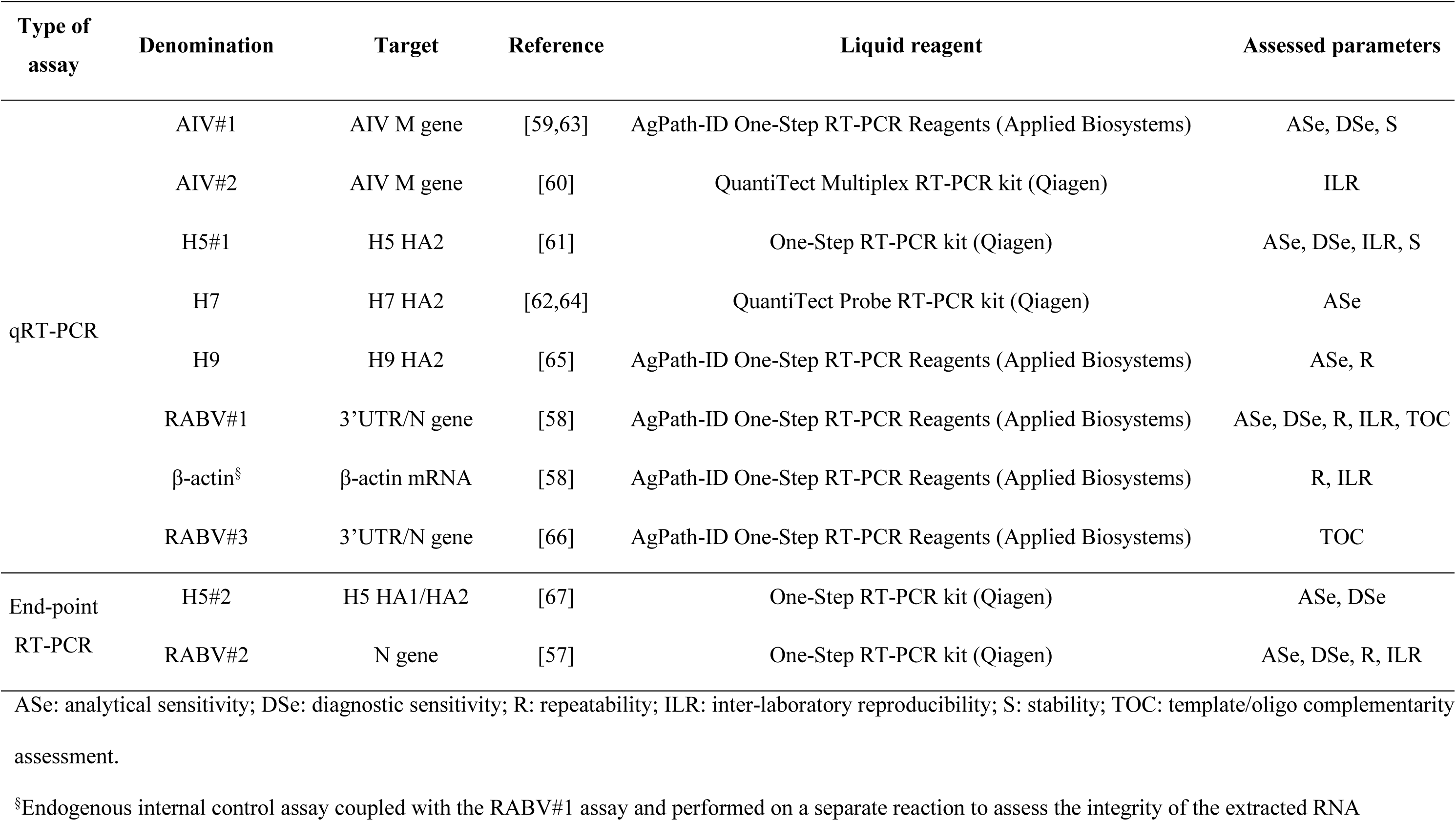
AIV and RABV molecular protocols employed to compare lyophilized and wet master mix reagents.

Reaction mix compositions and optimized amplification protocols are described in detail in S1 file. All qRT-PCR assays were performed on a CFX96 Touch Deep Well Real-Time PCR Detection System (Biorad) and analyzed with the CFX Maestro Software. End-point RT-PCR protocols were carried out on the S1000 Thermal Cycler (Biorad) and the amplification products were visualized with the QIAxcel Advanced System (Qiagen). Sanger sequencing downstream RT-PCR was performed using Genetic Analyzer 3500xL Dx (Applied Biosystems).

### Analytical sensitivity (ASe)

Analytical sensitivity was assessed testing ten-fold dilution series of synthetic RNA and viral isolates prepared in molecular-grade water and PBS with antibiotics/antimycotics (PBSa), respectively (Table 2). Synthetic RNA was produced from plasmid templates as described elsewhere [66]. Each dilution was tested in three replicas employing the selected lyophilized reagent, as well as the liquid master mixes for comparison. The limit of detection (LoD) was determined as the highest dilution at which all tested replicates yielded positive results. For end-point RT-PCR, positive results were confirmed by Sanger sequencing.

**Table 2.**
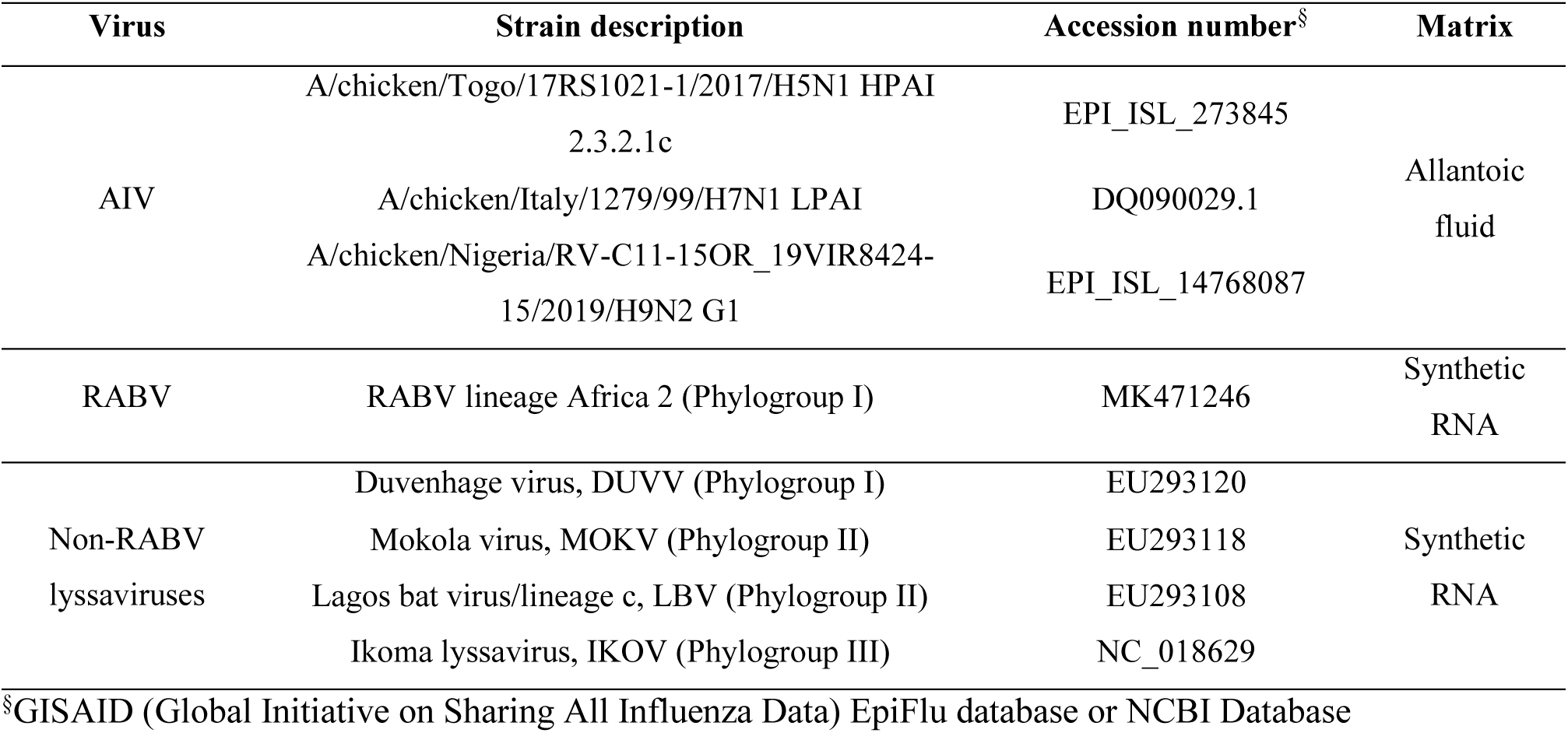
Samples used for analytical sensitivity.

### Repeatability within run and between days

Repeatability of Cq values obtained with the lyophilized reagent was assessed at two different concentrations of avian influenza A/chicken/Nigeria/RV-C11-15OR_19VIR8424-15/2019 (dilutions tested: 10^3.5^ and 10^1.5^ EID_50_/100 μl) and MOKV synthetic RNA (dilutions tested: 10^4^ and 10^2^ RNA copies/μl) employing the H9 and RABV#1 assays, respectively. Each dilution was tested in triplicate on two different days. The coefficient of variation (CV) of Cq values within run and between days was calculated to estimate repeatability.

### Diagnostic sensitivity (DSe)

Diagnostic sensitivity was assessed by testing avian influenza and rabies positive specimens retrieved from the IZSVe repository employing the freeze-dried reagent. They consisted of 24 bird samples (cloacal and oropharyngeal swabs, pool of organs) representing H5 HPAI viruses of clade 2.3.4.4b and 2.3.2.1c, and of 28 RABV infected brains representing the Cosmopolitan (Eastern Europe, Western European, and Africa 1 clades), Africa 2, Africa 3, Arctic-like and American indigenous lineages (S1 and S2 Tables).

Swab heads were cut, placed in 750-1000 µl PBSa and vortexed. Bird organs were diluted 1:4 w/v with PBSa and homogenized with the TissueLyser II (Qiagen) at 30 Hz for 3 min, while brain samples were homogenized in PBS (1:10 w/v) with glass rod. Homogenized tissues were subsequently clarified to obtain the supernatant. RNA was extracted from 100 µl of swab suspension or tissue supernatant. For avian influenza testing, the intype IC-RNA (Indical) was co-amplified with the viral genome (assay AIV#1) to ensure correct RNA extraction and verify the absence of PCR-inhibition. For RABV diagnostics, specimens conformity was verified by qRT-PCR targeting endogenous ß-actin mRNA [58]. For comparison, all samples were processed also with standard liquid reagents. Data were plotted using Prism 9.3.1 (GraphPad, San Diego, CA, USA), and the Spearman’s rank correlation coefficient (*r*) of the Cq obtained with the lyophilized and wet reagents was calculated. DSe and its exact binomial confidence interval (95% level) were assessed for each protocol using both the liquid and lyophilized reagents. Comparability of DSe between the freeze-dried and wet reagents was assessed by two-sample test of proportions. A *p*-value < 0.05 was considered significant.

### Stability verification of the lyophilized reagent

In order to verify the actual stability of the lyophilized reagent, we tested an H5N1 HPAI 2.3.2.1c isolate (A/chicken/Togo/17RS1021-1/2017) at high, medium and low concentrations (10^4.5^, 10^1.5^ and 10^0.5^ EID_50_/100 μl) with the AIV#1 and H5#1 assays after storing the freeze-dried kit under the following conditions: +4°C for 2 months, room temperature for 9 months and 30°C for 10 days. Storage at +4°C and room temperature not exceeding 9 months are guaranteed by the manufacturer, while reagent incubation at 30°C prior use was evaluated to mimic lapses that might occur during the reagents’ transportation in absence of controlled temperature. Samples were tested also with standard liquid reagents as a reference. The Mann-Whitney test was applied to verify any significant difference in terms of Cq values among the different storage conditions using Prism 9.3.1 (GraphPad, San Diego, CA, USA). A *p-*value < 0.05 was considered significant.

### Deployment of the lyophilized reagent through an inter-laboratory reproducibility study

An inter-laboratory reproducibility study was organized to deploy the lyophilized reagent for the molecular diagnosis of AI and rabies to CVLs of Sub-Saharan Africa. Four laboratories were invited to take part in the study, namely Laboratoire Central Vétérinaire (Mali), Laboratoire National d’Elevage et de Recherches Vétérinaires (Senegal), Laboratoire National Vétérinaire Yaoundé (Cameroon) and Laboratoire Central de Diagnostic Vétérinaire de Conakry (Guinea). At the time of the exercise, participating laboratories were routinely diagnosing AI and rabies through qRT-PCR and end-point RT-PCR, respectively. They all declared to have never applied qRT-PCR to diagnose rabies.

Participants were provided with (i) Standard Operating Procedures (SOPs) for AIV#2, H5#1, H5#2, RABV#1, β-actin and RABV#2 assays, to be performed with both lyophilized and liquid reagents, (ii) nucleic acid extraction kit, amplification kits and oligonucleotides necessary to perform the exercise, and (iii) panels of blind samples for avian influenza and rabies testing. Concomitantly to the shipment, an online meeting was arranged to provide appropriate instructions. At the end of the exercise, the laboratories were invited to submit results using a standardized data reporting form.

Test samples (RABV panel: *n* = 13; AIV panel: *n* = 12) comprising negative and positive specimens for the assessed targets were prepared using PrimeStore Molecular Transport Medium (MTM) (Longhorn Vaccines and Diagnostics, MD, USA) and appropriately blind-coded. Quality control tests (i.e. homogeneity and stability) were performed according to the ISO/IEC 17043:2010 and ISO 13528:2022, and using both liquid and freeze-dried reagents. Full panel compositions and expected results are detailed in S3 and S4 Tables.

### Data analyses of the inter-laboratory reproducibility study

#### For data analysis, laboratories were anonymized with unique letters (A-E)

First, we assessed whether the lyophilized reagent was able to correctly identify positive and negative samples for all the targets considered. In detail, we compared the results obtained by all five laboratories (four CVLs and the Reference Laboratory – RL, hosted by the IZSVe) with the freeze-dried reagent to the results obtained using the reference liquid master mixes by the RL. For each participant, we also evaluated the impact the lyophilized reagent could have on the target identification by comparing the results obtained at a laboratory level and between participants (freeze-dried Vs the liquid reagents). In both cases, the McNemar test was applied with matched pairs of dichotomous results (Positive, Negative) for all tested protocols but β-actin assay, for which the Stuart Maxwell test was used (k categories: Positive, Negative, Weakly positive). Cohen’s Kappa coefficient was also calculated and interpreted according to the Landis and Koch (1977) scale (< 0: no agreement; 0 – 0.20: slight agreement; 0.21 – 0.40: fair agreement; 0.41 – 0.60: moderate agreement; 0.61 – 0.80: substantial agreement; 0.81 – 0.99 almost perfect agreement; 1 perfect agreement).

Secondly, we assessed whether the differences between Cq values obtained with the lyophilized and liquid reagents were statistically significant through Bland-Altman plots. The normal distribution of Cq values was verified with the Shapiro-Wilk test. In case of normal distribution, the one sample t-test was performed to assess whether the mean bias was significantly different from zero, and the limits of agreement were calculated as mean bias ± 2 standard deviation. In case of non-normal Cq distribution, the limits of agreement were calculated as 2.5% and 97.5% percentiles of the differences distribution, and the significance of the median bias was tested using the Wilcoxon test. The mean or median bias, limits of agreement and the line of equality between liquid and lyophilized reagents were plotted into graphs.

Ultimately, we assessed the correct implementation of the qRT-PCR assay for rabies diagnosis at the four enrolled CVLs, employing RABV#1 and β-actin qRT-PCRs. In this context, statistical analysis was performed according to the number of ‘correct’ results, i.e. answers in agreement with the reference values assigned by the RL. All results were statistically analysed using either the McNemar or Stuart Maxwell tests, and the Cohen’s Kappa coefficient determination, as described above.

*P*-value < 0.05 was considered significant for all data analysis. Statistical software R version 4.2.1 was used to perform the statistical analyses.

## Results

### The Qscript lyo 1-step (Quantabio) was selected for analytical performance evaluation compared to standard liquid reagents

At the time of the study, we identified six different freeze-dried one-step qRT-PCR/end-point RT-PCR commercial products: the Qscript lyo 1-step (Quantabio, MA, USA), the Takyon Dry One-Step RT Probe MasterMix No Rox (Eurogentec, Belgium), the 5X CAPITAL qRT PCR Probe Master Mix lyophilized (Biotech rabbit, Germany), the SCRIPT RT-qPCR ProbesMaster Lyophilisate (Jena Science, Germany), the Lyophilized One Step qRT PCR (Biotechne SRL, Italy), and the ViPrimePLUS Lyophilized One Step qRT-PCR Master Mix (Vivanti, Malaysia). The main features of the above kits are detailed in S5 Table.

All multi-reaction kits requiring reconstitution of the entire lyophilized product prior reaction assembly were excluded due to the need to freeze the unused reagent after rehydration. In contrast, reagents pre-aliquoted as lyospheres in 200 μl tubes for single-reaction assembly were considered for further evaluation as they i) do not require storage at -20°C once reconstituted, ii) reduce the risk of cross-contamination, iii) allow higher repeatability by minimizing the effect of pipetting errors of the master mix, and iv) permit to use the tubes provided with no need to transfer the reaction mix, thus simplifying the reaction assembly. The Qscript lyo 1-step (Quantabio) and SCRIPT RT-qPCR ProbesMaster Lyophilisate (Jena) met such requirements and their ASe was preliminary assessed using the RABV#1 assay [58]. The Qscript lyo 1-step (Quantabio) proved to be more sensitive (data not shown) and was selected for subsequent evaluation of sensitivity.

For all qRT-PCR and end-point RT-PCR assays assessed and applied to AIV and RABV samples, the Qscript lyo 1-step kit showed the same LoD as the liquid master mixes. The internal control implemented for the AIV#1 assay provided conforming results in all cases. In contrast, lower analytical sensitivity was obtained for the H5#2 assay, which is based on an end-point RT-PCR reaction and yields an amplification products of 308 bp (decrease by 1 log) as well as for the RABV#1 and RABV#2 assays applied to non-RABV lyssaviruses (decrease up to 2 logs depending on the species) (Table 3).

**Table 3.** Analytical sensitivity (ASe) of the molecular protocols performed with the Qscript lyo 1-step kit and the liquid reagents.

| Assay | Sample | LoD (mean Cq ± SD) | |
| --- | --- | --- | --- |
|  |  | Liquid reagents | Lyophilized reagent |
| AIV#1 | H5N1 | 10<sup>0.5</sup> (35.8 ± 0.4) | 10<sup>0.5</sup> (35.0 ± 0.7) |
| H5#1 | H5N1 | 10<sup>0.5</sup> (36.0 ± 0.7) | 10<sup>0.5</sup> (35.6 ± 0.3) |
| H7 | H7N1 | 10<sup>0.5</sup> (35.2 ± 0.4) | 10<sup>0.5</sup> (37.8 ± 1.4) |
| H9 | H9N2 | 10<sup>0.5</sup> (31.7 ± 1.3) | 10<sup>0.5</sup> (34.1 ± 1.2) |
| H5#2 | H5N1 | **10<sup>-0.5</sup>** | **10<sup>0.5</sup>** |
| RABV#1 | RABV | 10 (33.8 ± 0.5) | 10 (33.4 ± 0.1) |
| | DUVV | **10<sup>2</sup>** (35.0 ± 0.5) | **10<sup>3</sup>** (30.9 ± 0.1) |
| | MOKV | 10<sup>2</sup> (33.8 ± 0.3) | 10<sup>2</sup> (34.2 ± 0.3) |
| | LBV | **10<sup>3</sup>** (33.4 ± 0.1) | **10<sup>5</sup>** (33.2 ± 0.3) |
| | IKOV | 10<sup>5</sup> (34.0 ± 0.1) | 10<sup>5</sup> (35.2 ± 0.2) |
| RABV#2 | RABV | 10<sup>2</sup> | 10<sup>2</sup> |
|  | DUVV | **10<sup>3</sup>** | **10<sup>4</sup>** |
|  | MOKV | **10<sup>3</sup>** | **10<sup>5</sup>** |
|  | LBV | 10<sup>7</sup> | 10<sup>7</sup> |
|  | IKOV | 10<sup>3</sup> | 10<sup>3</sup> |

LoD values are expressed as EID<sub>50</sub>/100 μl or RNA copies/ μl, according to the matrix employed. LoD values qualitatively different between the liquid and lyophilized reagents are highlighted **in bold**. For qRT-PCR assays, mean Cq and standard deviation values are reported in brackets.

The Qscript lyo 1-step (Quantabio) yielded repeatable results over the assays and virus concentration tested, with a CV of Cq values ≤ 0.04.

### The lyophilized reagent is suitable for clinical samples testing

All but one AIV-H5 positive samples representing different bird species, geographic origins and sample matrices were correctly identified by the AIV#1 and H5#1 assays employing the Qscript lyo 1-step (Quantabio), resulting in 96.55% DSe (Table 4). Most likely, sample 17RS115-4 was missed because of low viral load (S1 Table). The RABV#1 and RABV#2 assays employing the Qscript lyo 1-step (Table 4) correctly identified (100% DSe) all the tested RABV positive field samples. Overall, no significant differences were observed between lyophilized and liquid reagents, neither for avian influenza nor for rabies, with a high correlation of Cq values (*r* ≥ 0.904) (Figure 1).

**Fig1.**
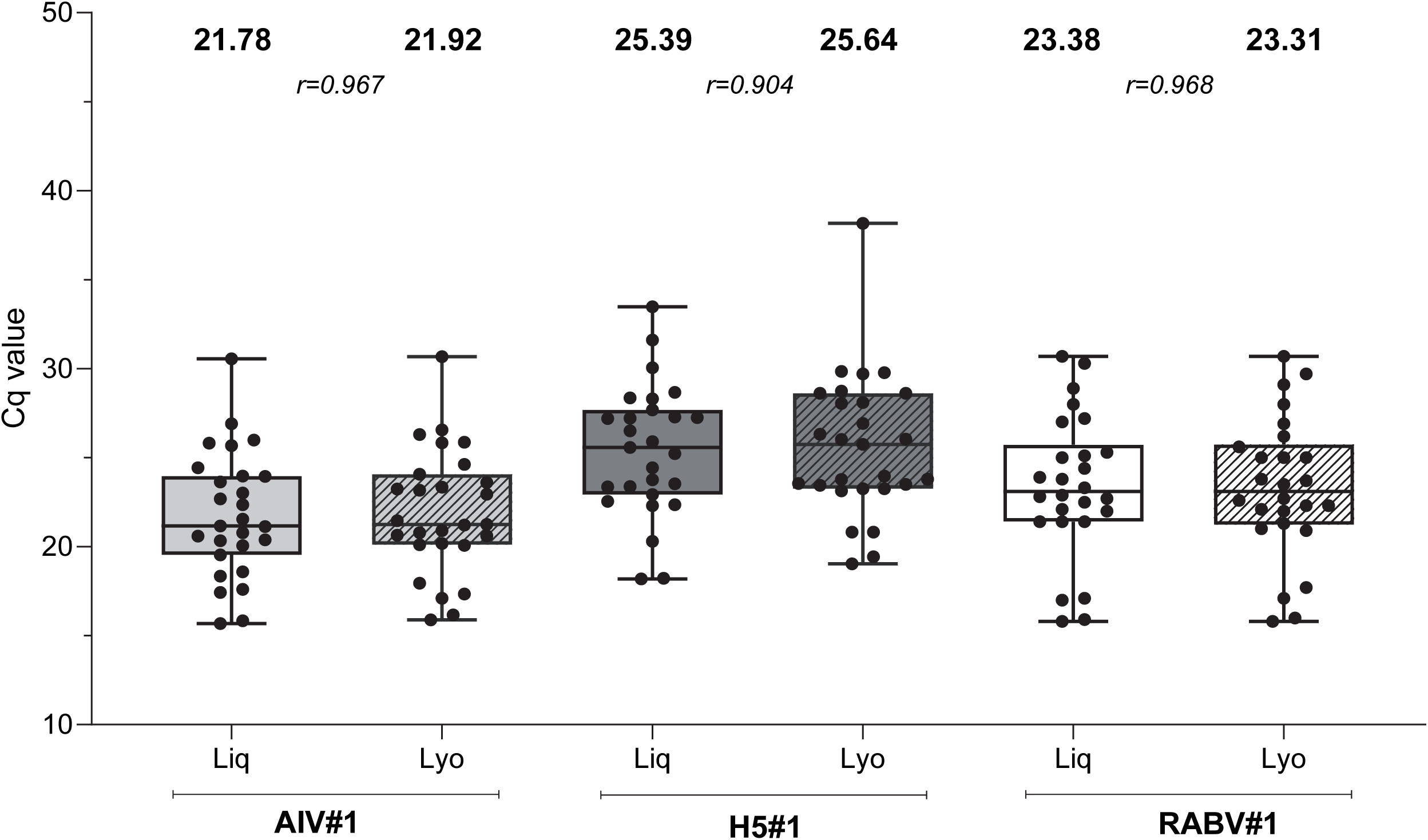
Diagnostic sensitivity of AIV#t, HS#t and RABV#t assays performed either with liquid and lyophilized reagents. Dots represent individual Cq values. Mean of Cq values for all samples at the different tested conditions (**in bold**) and Spearman’s rank correlation coefficient (*r*) between the liquid and the lyophilized reagents are reported. Liq: liquid reagent; Lyo: lyophilized reagent.

**Table 4.** Diagnostic Sensitivity (DSe) of molecular protocols performed with the Qscript lyo 1-step kit compared to the liquid reagents.

| Assay | DSe (95% CI) |  |
| --- | --- | --- |
|  | Liquid reagents | Lyophilized reagent |
| AIV#1 | 100 (88.05-100)* | 96.55 (82.23 – 99.91) |
| H5#1 | 96.55 (82.23 – 99.91) | 96.55 (82.23 – 99.91) |
| RABV#1 | 100 (87.23-100)* | 100 (87.23-100)* |
| RABV#2 | 100 (87.23-100)* | 100 (87.23-100)* |

DSe is expressed as percentage with 95% confidential interval; \* one-side – 97.5% CI.

### The performance of the lyophilized reagent might be affected by primer-template complementarity

After observing a decreased ASe of the Qscript lyo-1 step (Quantabio) compared to the liquid reagents when testing non-RABV lyssaviruses, we investigated more in depth the effect of oligonucleotide-template complementarity on the performance of the lyophilized reagent. To this aim, we applied the Qscript lyo-1 step (Quantabio) to the primers and probe from the (n) LN34 assay (RABV#3) which has been recently developed to improve coverage against divergent lyssaviruses [66]. LoD was re-assessed on serial dilutions of synthetic RNA from DUVV (phylogroup I), MOKV (phylogroup II) and IKOV (phylogroup III). Contrary to what observed for RABV#1, the ASe of RABV#3 [66] combined with the Qscript lyo-1 step (Quantabio) showed the same (DUVV) or improved (MOKV and IKOV) sensitivity compared to the liquid reagent (Figure 2, S6 Table). These results suggest that the performances of the Qscript lyo 1-step (Quantabio) might be affected by suboptimal primer-template complementarity.

**Fig.2.**
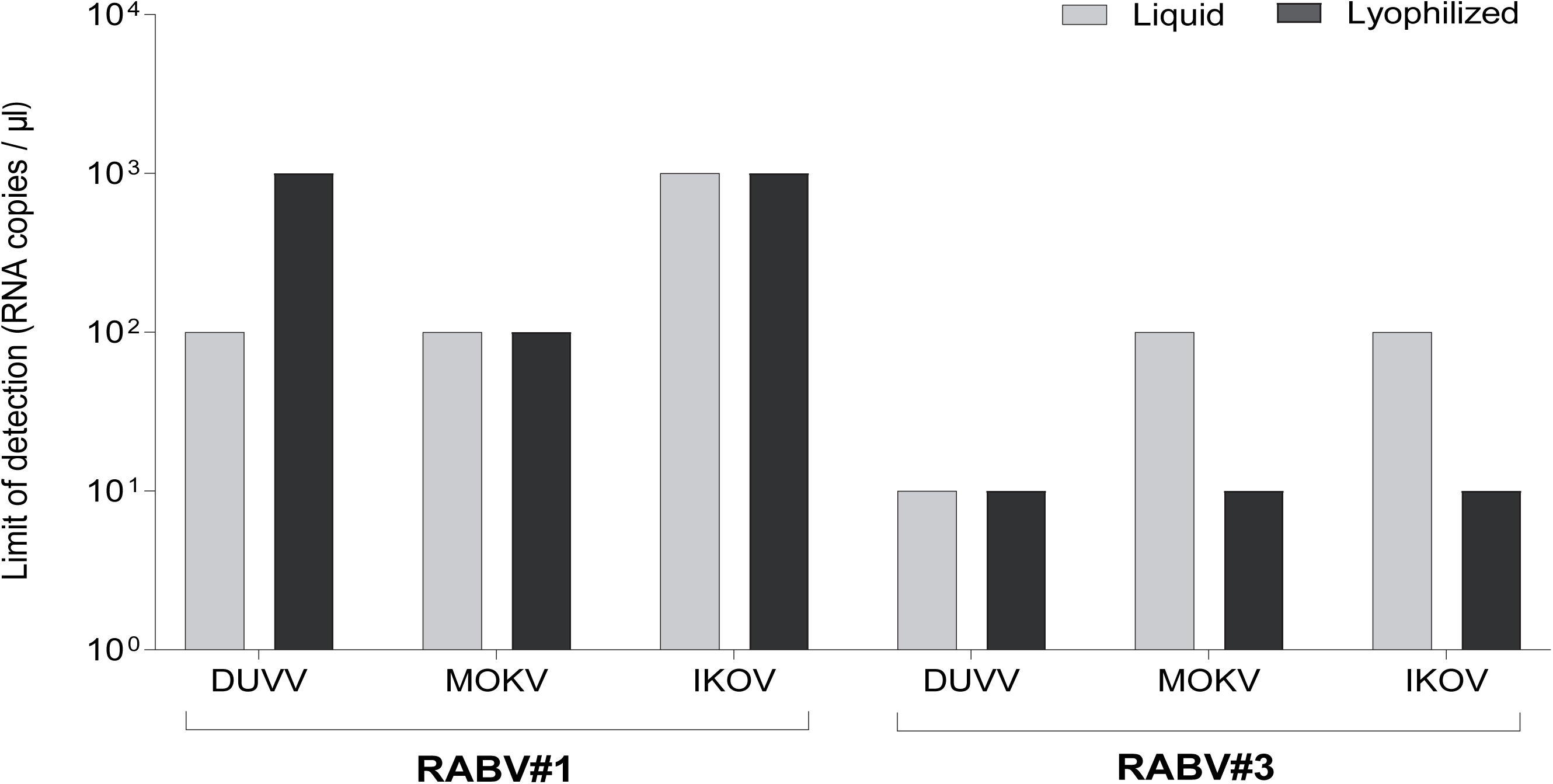
**Limit of detection of the RABV#l and the RABV#3 assays employing the lyophilized reagent Qscript lyo l-step (Quantabio) on serial dilutions of synthetic RNA from DUVV (phylogroup I), MOKV (phylogroup II) and IKOV (phylogroup III).**

### The lyophilized reagent shows optimal stability even when stored at 30°C for short periods

To evaluate the effect of environmental stressing conditions that might occur during reagents shipment or in the laboratory, the lyophilized kit was stored at different temperatures before use. Data obtained (Figure 3, S7 Table) indicate that the performance of the Qscript lyo 1-step (Quantabio) is comparable to the liquid reagents over the tested range of temperatures and incubation times, without any loss of sensitivity as assessed by the LoD. These comprise incubation for 10 days at 30°C, a scenario that might occur upon delayed reagents delivery or inadequate storage due to electricity blackouts at laboratory level. Notably, the use of the Qscript lyo 1-step (Quantabio) stored for 10 days at 30°C and for 9 months at room temperature showed slightly higher Cq values with respect to the lyophilized reagent stored under optimal conditions and to liquid reagents, although differences were not statistically significant (*p* > 0.05).

**Fig3.**
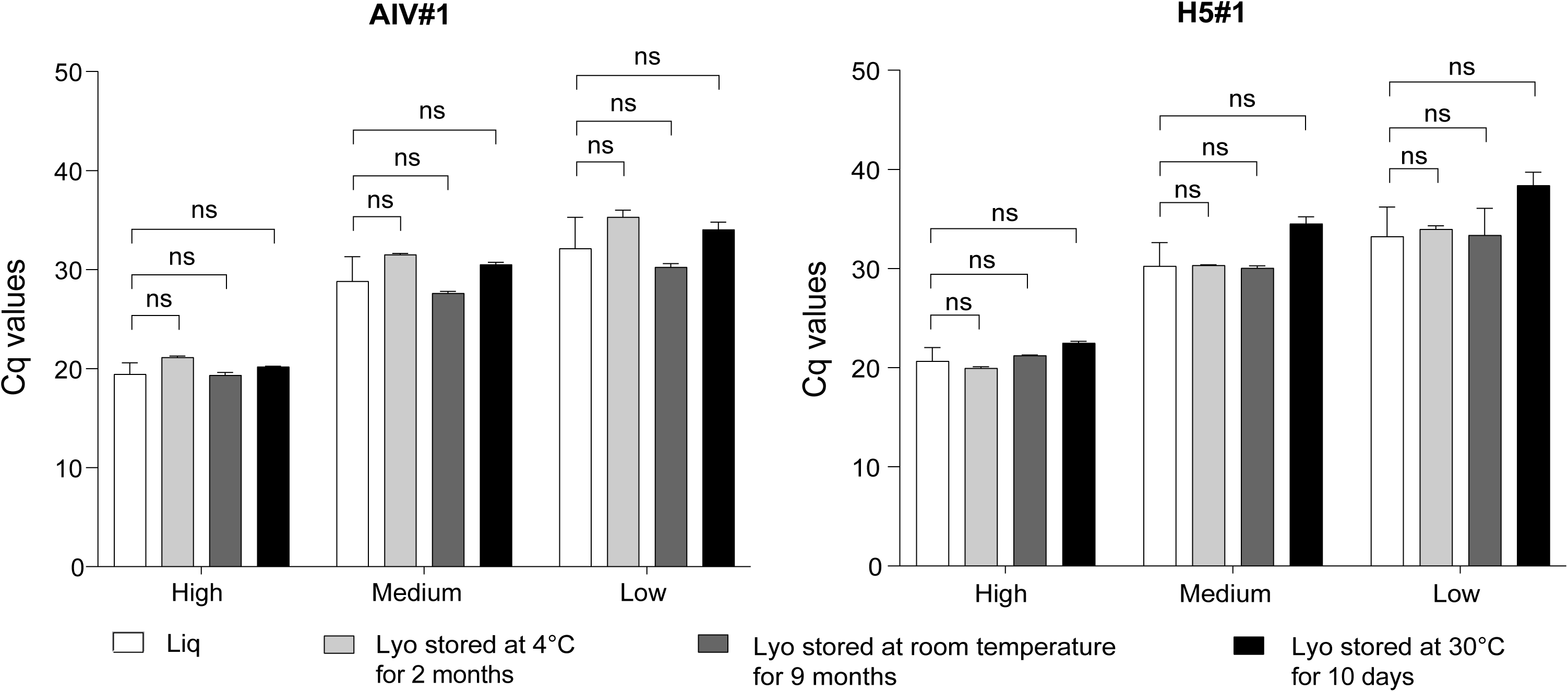
**Stability of the lyophilized kit under different storage conditions**. The Mann-Whitney test was applied to assess differences between mean Cq values of the reference (liquid reagents; Liq) and lyophilized reagent (Lyo) Qscript lyo 1-step. High, medium, and low concentrations of H5N1 (A/chicken/Togo/17RS1021-1/2017) were employed.

### The lyophilized reagent can be deployed in Sub-Saharan diagnostic laboratories for AIV and RABV molecular testing

The results obtained by all the laboratories taking part to the exercise and using the Qscript lyo 1-step kit (Quantabio) demonstrated perfect or almost perfect agreement with the reference results obtained by the RL (Cohen’s-Kappa 0.921–1) using liquid reagents, with similar Cq values (McNemar/Stuart Maxwell test p-value = 0.157 – 1) (Table 5).

**Table 5.** Agreement between the qualitative results of the Qscript Lyo-1 step kit (Quantabio) and the reference results.

| Assay | Cohen’s Kappa<sup>§</sup> | McNemar/Stuart Maxwell test<br>*p*-value |
| --- | --- | --- |
| AIV#2 | 0.963 | 0.317 |
| H5#1 | 0.931 | 0.157 |
| RABV#1 | 0.937 | 1 |
| β-actin | 0.921 | 0.223 |
| RABV#2 | 1 | n.a. |

Data from all participants obtained with the Qscript lyo 1-step kit (Quantabio) were compared to the reference results obtained by the RL that employed standard wet methods; n.a.: not applicable, § $p$ -value < 0.001.

When assessing the results obtained by each laboratory, statistical analyses revealed a perfect or almost perfect agreement between the Qscript lyo 1-step (Quantabio) and the liquid reagents for all the assays, except for β-actin in laboratory B, and RABV#1 and AIV#2 in laboratory D, which showed a substantial agreement (Table 6). The McNemar or Stuart Maxwell test demonstrated that there was no significant difference in the number of ‘correct’ results (Table 6).

**Table 6.** Agreement between Qscript Lyo-1 Step Kit (Quantabio) and liquid reagent qualitative results for each laboratory and overall.

| Assay | Test | Value | Laboratory |  |  |  |  | All labs |
| --- | --- | --- | --- | --- | --- | --- | --- | --- |
|  |  |  | A | B | C | D | E |  |
| AIV#2 | McNemar | *p*-value | n.a. | n.a. | n.a. | 0.317 | n.a. | 1 |
|  | Cohen's Kappa | *Kappa*<br>*p*-value | 1<br>< 0.001 | 1<br>< 0.001 | 1<br>< 0.001 | 0.800<br>0.002 | 1<br>< 0.001 | 0.925<br>< 0.001 |
| H5#1 | McNemar | *p*-value | 0.317 | 0.317 | n.a. | n.a. | n.a. | 0.157 |
|  | Cohen's Kappa | *Kappa*<br>*p*-value | 0.824<br>0.002 | 0.824<br>0.002 | 1<br>< 0.001 | 1<br>< 0.001 | 1<br>< 0.001 | 0.931<br>< 0.001 |
| RABV#1 | McNemar | *p*-value | n.a. | n.a. | n.a. | 1 | n.a. | 0.317 |
|  | Cohen's Kappa | *Kappa*<br>*p*-value | 1<br>< 0.001 | 1<br>< 0.001 | 1<br>< 0.001 | 0.675<br>0.007 | 1<br>< 0.001 | 0.872<br>< 0.001 |
| β-actin | Stuart Maxwell | *p*-value | n.a. | 0.367 | 0.606 | n.a. | n.a. | 0.007 |
|  | Cohen's Kappa | *Kappa*<br>*p*-value | 1<br>< 0.001 | 0.745<br>< 0.001 | 0.851<br>< 0.001 | 1<br>< 0.001 | 1<br>< 0.001 | 0.723<br>< 0.001 |
| RABV#2 | McNemar | *p*-value | n.a. | n.a. | n.a. | n.p. | n.a. | n.a. |
|  | Cohen's Kappa | *Kappa*<br>*p*-value | 1<br>< 0.001 | 1<br>< 0.001 | 1<br>< 0.001 | n.p.<br>n.p. | 1<br>< 0.001 | 1<br>< 0.001 |

n.a: not applicable; n.p.: analysis not performed.

Perfect or almost perfect agreement between aggregated data obtained from all laboratories with the lyophilized reagent compared to liquid master mixes was calculated for all assays but β-actin, which showed a substantial agreement (Cohen’s-Kappa 0.723; Stuart Maxwell test p-value = 0.007). The Bland Altman plots revealed no significant differences in terms of Cq values between the Qscript lyo 1-step kit (Quantabio) and the liquid reagents for AIV#1 and RABV#1 assays, while Cq values decreased for the lyophilized reagent when applying the H5#1 assays, meaning that freeze-dried and wet master mixes are interchangeable. In contrast, an increase in Cq values was observed for the β-actin assay (Figure 4).

**Fig4.**
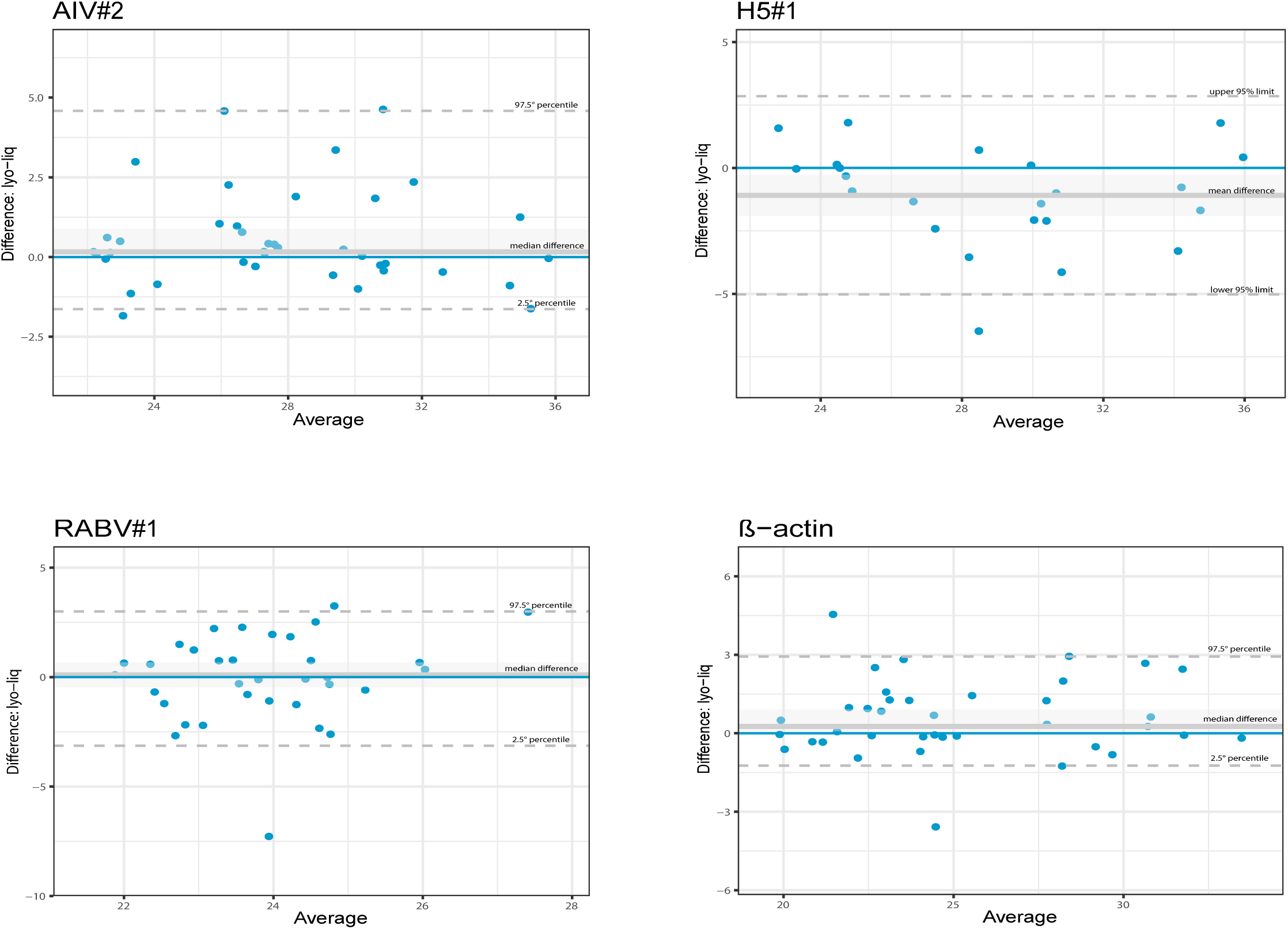
The Bland Altman plots showing differences between Ct values of liquid reagents vs the AIV#2, HS#l, RABV#l, and β-actin rRT-PCR protocols run with Qscript lyo l-step. The blue lines represent perfect agreement. The grey lines represent the mean or median bias (according to the satisfaction or not of normality requirement) between the lyophilized ("lyo") and liquid ("liq") reagents. The grey dotted lines represent the upper and lower limits of agreement. **A) AIV#2:** The differences between the two methods are within the -1.630 and 4.582 range, and the two methods may be used interchangeably. The median bias of 0.165 was not significantly different from zero (Wilcoxon test p-value = 0.082, the line of equality is within the confidence interval of the median difference); **B) HS#l:** The differences between the two methods are within the -5.024 and 2.850 range. The mean bias -1.087 was significantly different from zero (T-test test p-value = 0.0165, the line of equality is not within the confidence interval of the mean difference). Bland Altman’s diagram shows a substantial systematic difference between the two methods, resulting in better performances of the lyophilized reagent; **C) RABV#l:** The differences between the two methods are within the -3.130 and 2.998 range, the two methods can be used interchangeably. The median bias of 0.103 was not significantly different from zero (Wilcoxon test p-value = 0.774, the line of equality is within the confidence interval of the median difference); **D) β-actin:** The differences between the two methods are within the -1.240 and 2.934 range. The mean bias -0.260 was significantly differed from zero (Wilcoxon test p-value = 0.027, the line of equality is not within the confidence interval of the median difference). The Bland Altman’s diagram shows a substantial systematic difference between the two methods leading to worse performances with lyophilized reagents.

### Rabies qRT-PCR was successfully implemented in Sub-Saharan CVLs

As the CVLs had declared no prior experience in implementing RABV#1, and more in general qRT-PCR assays to diagnose rabies, we compared the results obtained using the reference liquid master mix at their laboratories with those from the RL. Perfect or almost perfect agreement was observed between the RL and the enrolled CVLs with only one exception, for which substantial/fair agreement was achieved for the RABV#1/β-actin assays, respectively (Table 7).

**Table 7.** Implementation of the qRT-PCR for rabies diagnosis in Sub-Saharan laboratories.

| Assay | Test | Value | Laboratory |  |  |  | All labs |
| --- | --- | --- | --- | --- | --- | --- | --- |
|  |  |  | A | B | C | D |  |
| RABV#1 | McNemar | *p-value* | 0.157 | n.a. | n.a. | n.a. | 0.157 |
|  | Cohen's Kappa | *Kappa* | 0.698 | 1 | 1 | 1 | 0.920 |
|  |  | *p-value* | 0.004 | < 0.001 | < 0.001 | < 0.001 | < 0.001 |
| β-actin | Stuart Maxwell | *p-value* | n.a. | n.a. | n.a. | n.a. | 0.030 |
|  | Cohen's Kappa | *Kappa* | 0.435 | 0.851 | 1 | 0.851 | 0.753 |
|  |  | *p-value* | 0.002 | < 0.001 | < 0.001 | < 0.001 | < 0.001 |

n.a.: not applicable.

## Discussion

Unreliable electricity provision as well as supply chain issues under refrigeration or frosting are recognized among the most significant factors that impact the ability to perform accurate lab-work in Sub-Saharan Africa. In these scenarios lyophilized reagents for molecular testing appear as effective solutions to overcome such constrains, as they are developed to combine higher stability and temperature resilience, simpler transportation and storage, space efficiency, ease of use, higher repeatability and reproducibility and, ultimately, cost effectiveness.

Among the different freeze-dried formulations commercially available, we selected the Qscript lyo 1-step (Quantabio) for its single-reaction format that offers numerous advantages in terms of conservation, reaction assembly, reliability and performance. A previous work has preliminarily demonstrated the utility of this product for AIV screening by qRT-PCR [56]. However, the current study provides the first comprehensive evaluation of its functionality throughout the entire diagnostic workflow, starting from the nucleic acids extraction to AIV subtyping by both qRT-PCR/RT-PCR and Sanger sequencing, and has further extended its validation to rabies diagnosis. Importantly, our data indicate that the replacement of liquid reagents with the Qscript lyo 1-step (Quantabio) in a wide range of qRT-PCR and end-point protocols produces repeatable results without significantly affecting sensitivity for the majority of the targets evaluated, as assessed by LoD determination and diagnostic performance on positive clinical samples. However, it is worth noting that the end-point RT-PCR H5#2 yielded a LoD improved by 1 log when using the wet reagents. It can be assumed that the Qscript lyo 1-step (Quantabio) sensitivity slightly decreases when longer targets are amplified, as a result of suboptimal reaction kinetics that are specific to each protocol and warrant evaluation on a case by case basis. Another condition that most likely affected the functionality of the lyophilized reagent is the suboptimal oligonucleotides complementarity [56,68]. Indeed, we observed a decrease in sensitivity when we applied the Qscript lyo 1-step kit (Quantabio) to DUVV and LBV with the RABV#1 assay. Notably, the detection of divergent lyssaviruses was restored with the RABV#3 assay, a method specifically designed to widen assay inclusivity and sensitivity by enhancement of the primer-template complementarity [66]. Although the authors acknowledge that further evaluation is necessary to recommend the RABV#3 in place of RABV#1 assay using the lyophilized reagent, our results support the hypothesis that the Qscript lyo 1-step kit (Quantabio) performance depends, at least in part, on oligonucleotides complementarity towards the target region and underlines the need for a careful evaluation of the diagnostic method well before implementing the use of lyophilized reagents.

Importantly, to accomplish the deployment of AI and RABV diagnostic assays based on the use of the lyophilized reagent, our validation scheme included an inter-laboratory reproducibility exercise that engaged four CVLs located in Sub-Saharan Africa. Data submitted by the participants confirmed a perfect or almost perfect agreement with the expected results and demonstrated that the implementation of the lyophilized reagent in place of liquid formulations allowed the correct identification of AIV and RABV in most cases, resulting in highly reproducible results among different laboratories. Considering rabies diagnosis, it is worth noting that the four CVLs had no prior experience with the RABV#1 assay or with any other qRT-PCR protocol for the diagnosis of rabies, but were anyway able to successfully implement such techniques at their facilities. This lack of experience may partly explain the suboptimal results observed for the β-actin assay using both the liquid and the lyophilized master mixes. Although improvable, the successful acquisition of qRT-PCR protocol by the four participating CVLs represents a significant achievement for surveillance and confirms that molecular assays can be easily implemented in tropical regions [37], specifically in veterinary laboratories that have been trained and supported to enable early diagnosis of other priority animal diseases [69–72]. Notably, the participating laboratories reported no false positives during the inter-laboratory exercise, further confirming the ease of use and reliability of the lyophilized reagent format. Furthermore, the simultaneous application of wet reagents throughout the trial allowed a quantitative evaluation of the Qscript lyo 1-step kit (Quantabio) performance applied to qRT-PCR assays, which revealed an irrelevant shift of Cq values compared to liquid reagents (as assessed by intra- and inter-laboratory comparisons) and demonstrated that the freeze-dried and wet reagents can be used interchangeably.

Another key feature that further confirms the feasibility of using the lyophilized reagent in Sub-Saharan regions is its thermal stability. According to the manufacturer’s recommendations, the Qscript lyo 1-step kit (Quantabio) can be stored at room temperature (21-23°C) for up to 9 months. However, temperature can vary during transportation, and storage is barely controlled in tropical and subtropical areas. We therefore challenged the lyophilized kit by incubation at 30°C for 10 days to mimic the absence of cold-chain maintenance [37]. The Qscript lyo 1-step kit (Quantabio) proved to be stable under such conditions, and at the same time resilient in case of electricity discontinuation at laboratory facilities or delayed intercontinental parcel delivery at uncontrolled temperatures.

In conclusion, our study demonstrates that lyophilized reagents can replace liquid formulations for AI and RABV molecular diagnosis, and provide reliable results under real Sub-Saharan laboratory settings. The successful application of the Qscript lyo 1-step kit (Quantabio) on a variety of assays and targets suggests that its use has the potential to be extended to the molecular diagnosis of other relevant veterinary and zoonotic diseases in remote areas characterized by poor transportation and storage infrastructures. However, on a larger scale, the implementation of lyophilized reagents requires accurate optimisation of the amplification conditions and a throughout performance evaluation of the selected protocols.

## Acknowledgments/Fundings

Most of the activities were supported by UN-FAO with funding from the US Agency for International Development (USAID) under the Global Health Security Agenda (GHSA)/ Global Health Security Project (GHSP). The content of this article is the responsibility of the author(s) and does not necessarily reflect the views of UN-FAO, USAID, or the US Government.

## Author Contributions

**Conceptualization:** Paola De Benedictis, Isabella Monne, Valentina Panzarin

**Data curation:** Petra Drzewnioková, Irene Brian, Marzia Mancin, Andrea Fortin, Valentina Panzarin.

**Formal analysis:** Petra Drzewnioková, Irene Brian, Marzia Mancin, Andrea Fortin.

**Funding Acquisition:** Morgane Gourlaouen, Angélique Angot, Mamadou Niang, Baba Soumare, Paola De Benedictis, Isabella Monne, Valentina Panzarin.

**Investigation:** Petra Drzewnioková, Irene Brian, Marzia Mancin, Andrea Fortin, Isaac Dah, Kouramoudou Berete, Adama Diakite, Fatou Tall Lo, Valeria D’Amico, Viviana Valastro.

**Methodology:** Petra Drzewnioková, Marzia Mancin, Paola De Benedictis, Isabella Monne, Valentina Panzarin.

**Project Administration:** Paola De Benedictis, Isabella Monne, Valentina Panzarin.

**Resources:** Morgane Gourlaouen, Angélique Angot, Mamadou Niang, Clement Meseko, Emilie Go-Maro, Viviana Valastro, Baba Soumare, Paola De Benedictis, Isabella Monne, Valentina Panzarin.

**Software:** Marzia Mancin.

**Supervision:** Paola De Benedictis, Isabella Monne, Valentina Panzarin.

**Validation:** Petra Drzewnioková, Irene Brian, Marzia Mancin, Andrea Fortin, Valentina Panzarin. **Visualization:** Petra Drzewnioková, Irene Brian, Marzia Mancin, Paola De Benedictis, Isabella Monne, Valentina Panzarin.

**Writing – Original Draft Preparation:** Petra Drzewnioková, Irene Brian, Marzia Mancin, Paola De Benedictis, Isabella Monne, Valentina Panzarin.

**Writing– Review & Editing:** All authors.

